# Mobile steady-state evoked potential recording: dissociable neural effects of real-world navigation and visual stimulation

**DOI:** 10.1101/705095

**Authors:** James Dowsett, Marianne Dieterich, Paul C.J. Taylor

## Abstract

**Background:** The ability to record brain activity in humans during movement, and in real world environments, is an important step towards understanding cognition. Electroencephalography (EEG) is well suited to mobile applications but suffers from the problem of artefacts introduced into the signal during movement. Steady state visually evoked potentials (SSVEPs) give an excellent signal-to-noise ratio and averaging a sufficient number of trials will eventually remove any noise not phase locked to the visual flicker.

**New Method:** Here we present a method for producing SSVEPs of real world environments using modified LCD shutter glasses, which are commonly used for 3D TV, by adapting the lens to flicker at neurophysiologically relevant frequencies, in this case the alpha band. Participants viewed a room whilst standing and walking. Either the left or right side of the room was illuminated, to test if it is possible to recover the resulting SSVEPs when walking, as well as to probe the effect of walking on neural activity. Additionally, by using a signal generator to produce “simulated SSVEPs” on the scalp we can demonstrate that this method is able to accurately recover evoked neural responses during walking.

**Results:** The amplitude of SSVEPs over right parietal cortex was reduced by walking. This finding is in line with converging evidence that visual-vestibular integration involves cortical lateralization with the right hemisphere being dominant in right handers. Furthermore, the waveform and phase of the SSVEPs is highly preserved between walking and standing, but was nevertheless sensitive to whether visual stimulation was presented to the left or right visual hemifield.

**Conclusions:** This method allows probing neural responses at a wide range of frequencies, during natural movements within real environments.

## Introduction

Most methods used to study human brain activity, such as fMRI, MEG and conventional EEG, require the head and body to be in a fixed position. In the case of EEG and MEG, it is usually also necessary for the eyes to fixate, to prevent eye movement artefacts. Developing methods for studying brain activity whilst moving and looking around natural environments is arguably an important step towards elucidating human cognition (Gramann et al., 2014; Boto et al., 2018). The main aim of the current study is to develop a method to create evoked responses to real world visual scenes during mobile EEG when the participant is moving. Specifically we demonstrate a method which can provide robust neural signals during whole body locomotion, which can also be applied during head and eye movements.

In the current study we were particularly interested in the lateralization of neural oscillations, as visual-vestibular interaction is thought to rely on lateralization within the vestibular system driven by the non-dominant hemisphere, i.e. right hemisphere in right handers (Dieterich et al., 2003; Dieterich & Brandt, 2015, 2018). Such lateralization has been observed in humans during virtual navigation (Jacobs et al., 2010), but to date has not been observed in EEG during natural movement whilst observing real visual scenes.

Mobile EEG has made significant progress in recent years due to the decrease in the size and cost of amplifiers, allowing small unobtrusive recording devices which can be worn during everyday activities. Reliable EEG data suitable for mobile brain-computer interfaces has been demonstrated (Debener, Minow, Emkes, Gandras, & de Vos, 2012). However, the problem of motion artefacts has always proved challenging. The most popular strategy is to use independent component analysis (ICA), or some variation thereof, to separate out movement and muscle artefacts from meaningful neural signal. A practical limitation of ICA is that it requires a large number of electrodes, typically 64. Although mobile EEG systems with large numbers of electrodes do exist the setup time is a limitation; recording EEG in real world situations and in patient studies would benefit from faster setups.

A further problem with ICA is the potential to introduce additional sources of variance as different algorithms can give different results (Pontifex, Gwizdala, Parks, Billinger, & Brunner, 2017). Some researchers have used gait based movement artefact template subtraction to recover steady state visual evoked potentials (SSVEPs) and the P300 component of ERPs during movement (Kim & Jo, 2015). More recently, attempts to remove motion artefacts from mobile EEG using the electrocardiogram (ECG) signal as a reference signal have been able to reduce, but not completely remove, the movement-related artefacts (Butkeviciute et al., 2019).

SSVEPs are a method which typically involves various elements on a display (monitor) flickering at one or more frequencies which can be measured in the EEG signal (Norcia, Appelbaum, Ales, Cottereau, & Rossion, 2015; Haegens & Zion Golumbic, 2018; Vialatte et al., 2010). SSVEPs are popular partly because of the excellent signal-to-noise ratio achieved by averaging many segments of data. In theory, a large enough number of segments being averaged will remove any signal which is not exactly at the flicker frequency (or a higher harmonic) in the process of averaging. A few studies to date have combined SSVEPs with mobile EEG and demonstrated that the flicker signal can in principle be recovered (Lin, Wang, Wei, & Jung, 2014).

Liquid crystal display active 3D glasses (LCD glasses) are commonly used at particular frequencies for 3D TV. Here we adapt this technology to flicker at other frequencies, and with both eyes receiving the same flicker phase. This allows the generation of SSVEPs from whatever the participant is looking at, at frequencies not limited to multiples of a screen refresh rate as with SSVEP paradigms using regular computer monitors.

Here we manipulated the visual scene by selectively illuminating the left or right half of the room in which the participants were walking, and therefore selectively stimulating left or right hemifield. This was done to provide two visual conditions, to test whether the difference between the resulting SSVEPs in the standing condition could be reproduced in the walking condition, as well as to investigate the relative involvement of hemispheric lateralisation. The hypothesis was a different response to left hemifield illumination for standing and walking. This would be consistent with the additional processing demands of visual-vestibular interaction during walking in the right hemisphere (for right handed participants). In a separate condition we used a signal generator to produce “simulated SSVEPs” on the scalp as a control to demonstrate that this method is able to accurately recover SSVEPs during walking.

## Methods

### Experimental design

Ten normal right handed participants were recruited (six female, mean age 25 years, S.D. 2.46, mean Edinburgh handedness score 82.5). The study was approved by the local ethics committee (LMU Medical Faculty).

The experiment was conducted in a large empty room (7 × 5 m) with two facing white walls. Two black fixation crosses (7.5 × 8 cm) were attached in the centre of each wall at the eye height of the participant. The fixation crosses were illuminated by two spotlights, positioned immediately below each of the crosses and centred on them. Other than the two spotlights there were no other sources of light in the room. The spotlights were adapted with rectangular cardboard side-shutters which could be adjusted to block the light on either the left or right side. This resulted in only one side of the room being illuminated with a sharp vertical shadow centred on the fixation cross. In this way the visual stimulus (the illumination) was presented either to the left or right visual field. The main experiment consisted of four blocks of flicker, followed by a control experiment with a signal generator instead of the flicker.

**Figure 1:**
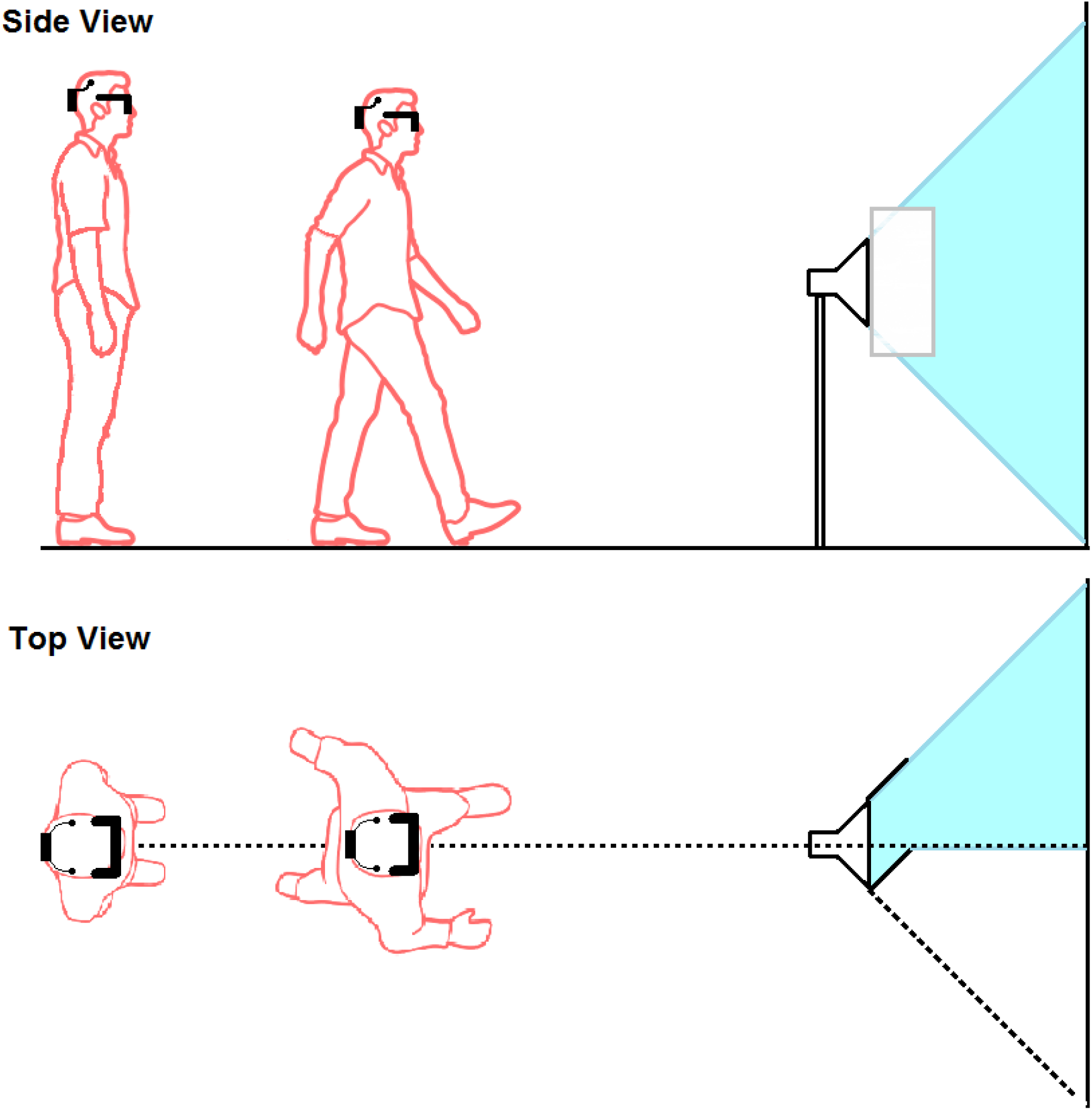
Schematic of experimental procedure, showing a trial with the visual stimulus in the left visual field.

### Main Experiment

Blocks alternated between walking and standing. Each block was 5 minutes long. During the first two blocks the side-shutters on the spotlights were adjusted such that one side of the room relative to the participant was illuminated, and for the second two blocks the lighting was adjusted so the other side was illuminated, e.g. if the left hemifield was illuminated for the participant for the first two blocks, the right hemifield was illuminated for the second two. Order of left and right hemifield was counter-balanced across participants.

### Participants walking

On walking blocks participants were instructed to fixate the cross on the far wall and to walk at their preferred walking speed towards it: mean stride frequency, as calculated through FFT analysis of the accelerometer data, was 1.45 Hz (max: 1.7 Hz, min: 0.8 Hz). When participants reached a distance of approximately one metre from the far wall, they were to stop, turn, fixate on the fixation cross on the other wall, pause for one second, and then to continue to walk towards the other wall, again while fixating. Walking distance across the room was approximately 5 meters and participants on average walked the length of the room 30.4 times per 5 minute block. Average time to walk the length of the room was 5.8 seconds giving an average walking speed of approximately 0.87 meters per second (standard deviation 0.15 m/s). Although this is slower than typical preferred walking speed of 1.42 m/s (Browning, Baker, Herron, & Kram, 2006), this is most likely due to the limited distance of 5 meters and the necessity to decelerate before stopping at the opposite wall.

### Participants standing

For the standing blocks the participants were instructed to stand still in the initial starting position, where the walking began, and fixate on the cross at the far end of the room. Every 20 seconds the experimenter would instruct the participant to take one step forward. When the participant reached one metre from the far wall, the participant was instructed to turn, fixate on the cross on the opposite wall, and then continue stepping towards it once every 20 seconds when prompted. This resulted in each participant moving the length of the room and back during the 5 minute block. This was done to match the visual input across the two conditions: each standing and walking block contained approximately the same amount of time spent at different distances to each wall, the remaining difference between conditions being the motion.

### EEG recording

EEG was recorded throughout the experiment at 1000 Hz with a LiveAmp 8 channel mobile EEG system (Brain Products, Munich Germany) which includes a 3-axis accelerometer in the amplifier, and was attached to the back of the EEG cap. Scalp electrodes (Ag/AgCl ring electrodes, BrainCap, Brain Products, Munich, Germany) were positioned according to the 10-10 system at O1, O2, P3, P4, P7 and P8. Impedance was kept below 10 kΩ and data was recorded with a sampling rate of 1000 Hz. The online reference was the right ear lobe, with a second reference attached to the left ear, and data was re-referenced offline to the average of the two. The ground electrode was positioned at electrode position Fpz and an EOG electrode was positioned below the right eye to record eye blinks. Setup time with this number of electrodes was approximately 5 minutes.

### LCD shutter glasses

For the main experiment (first four blocks) participants were wearing custom built adapted LCD glasses (SainSonic, Texas, US), controlled by a microcontroller (Arduino Uno, Scarmagno, Italy). When a voltage is applied across the layer of liquid crystal within the glass it becomes darker. Under their intended use the left and right eye rapidly and alternately become opaque (typically 120Hz), synchronized to the presentation of two images on the screen, each taken from slightly different viewpoints such that only one is visible to each eye, to create the illusion of one 3D image. Here the glasses were set to flicker at 10 Hz by eliciting a train of 5 volt pulses lasting 50 ms, which darkened the glass for both eyes; each pulse was followed by 50 ms with no voltage during which the glass was transparent, resulting in a 100 ms cycle. During the darkened part of the cycle the glass is not completely opaque but rather provides a low light view similar to wearing sunglasses. The pulse signal controlling the glasses was split and used as a trigger into the EEG amplifier for later segmentation of the data. The glasses were adapted with black cardboard frames occluding far peripheral vision beyond the frames, limiting the visual angle to 75 degrees in the horizontal plane.

### Control experiment

A common problem with any movement artefact removal strategy is that the use of real data precludes the ability to know the “true” uncontaminated EEG. Movement artefacts and movement related brain activity are not independent and are likely to co-vary. To test the efficacy of our method we used a signal generator, attached to the head of the participants, to deliver a weak voltage across the scalp. The voltage was adjusted such that the resulting signal in the surrounding electrodes, when averaged, was approximately the same size as the visually evoked potential of interest. By creating simulated SSVEPs in this way the number of segments required to average out the additional noise introduced by walking can be estimated by randomly selecting subsets of the available segments to create evoked potentials, and repeating this many times for different numbers of segments from the walking condition and comparing the result to the standing condition, which is uncorrupted by motion artefacts. Using this method we can gauge the signal-to-noise ratio expected from the addition of movement artefacts as we know the signals are identical. This allows the testing of a range of different walking speeds and gaits across participants, and therefore allows us to determine the reliability of the results.

The signal generator condition consisted of one walking block and one standing block. For these two blocks participants performed the same task with 5 minutes walking followed by 5 minutes standing, but without the glasses, and with both left and right hemifield illuminated. Instead, a signal generator (SIGGI 2, Easycap, Germany) was used to create electrical signals within the normal voltage range of typical EEG. The artificial signal (1mV, 10 Hz square wave) was applied throughout the experiment to EEG electrodes which were then attached to the electrodes at positions P7 and P8 using double-sided adhesive rings (which are typically used to attach electrodes on the skin), and filled with conductive gel. Pilot data indicated that this voltage applied to P7 and P8 produced a signal in electrodes O1, O2, P3 and P4, which was approximately the same peak-to-peak amplitude as SSVEPs resulting from wearing the LCD glasses flickering at 10 Hz. Subsequent inspection of the data confirmed this to be the case across participants: with a mean peak-to-peak amplitude of between 3 and 7 µV for all SSVEPs and “simulated SSVEPs” (Figure 3).

### Data analysis

For the walking blocks, data was rejected during the period when participants were stopping or turning. This was done by selecting 500 ms segments of data from the vertical axis channel of the accelerometer, centred on each trigger time. The vertical axis gives a clear spike with each footfall during walking (Figure 2, left panel); segments with a range of less than 50 m*g* vertical indicated the participant was not walking (i.e. was standing or turning); 500 ms was used to ensure that enough time was included in the accelerometer segment to capture this steep spike for participants with slower walking frequency. For the standing blocks, triggers were rejected if there was any movement in the vertical accelerometer above 20 mɡ, which rejected data during the time when the participants were stepping forward. Additionally data were visually inspected to ensure no triggers remained during standing and turning in the walking block. Finally, any data segment with a range of greater than 100 μV in the EOG channel was rejected to remove eye blinks (eye blinks were clearly visible in the data even when walking and small stabilizing eye movements due to fixating while walking were not visible in the raw EOG). After all segments were rejected there was an average of 1892 (S.D. = 418.2) segments per participant in the walking condition with flicker, 1924 (S.D. = 286.9) in the walking signal generator condition, 2337 (S.D. = 266.3) in the standing condition with flicker, and 2374 (S.D. = 337.5) in the standing signal generator condition.

**Figure 2:**
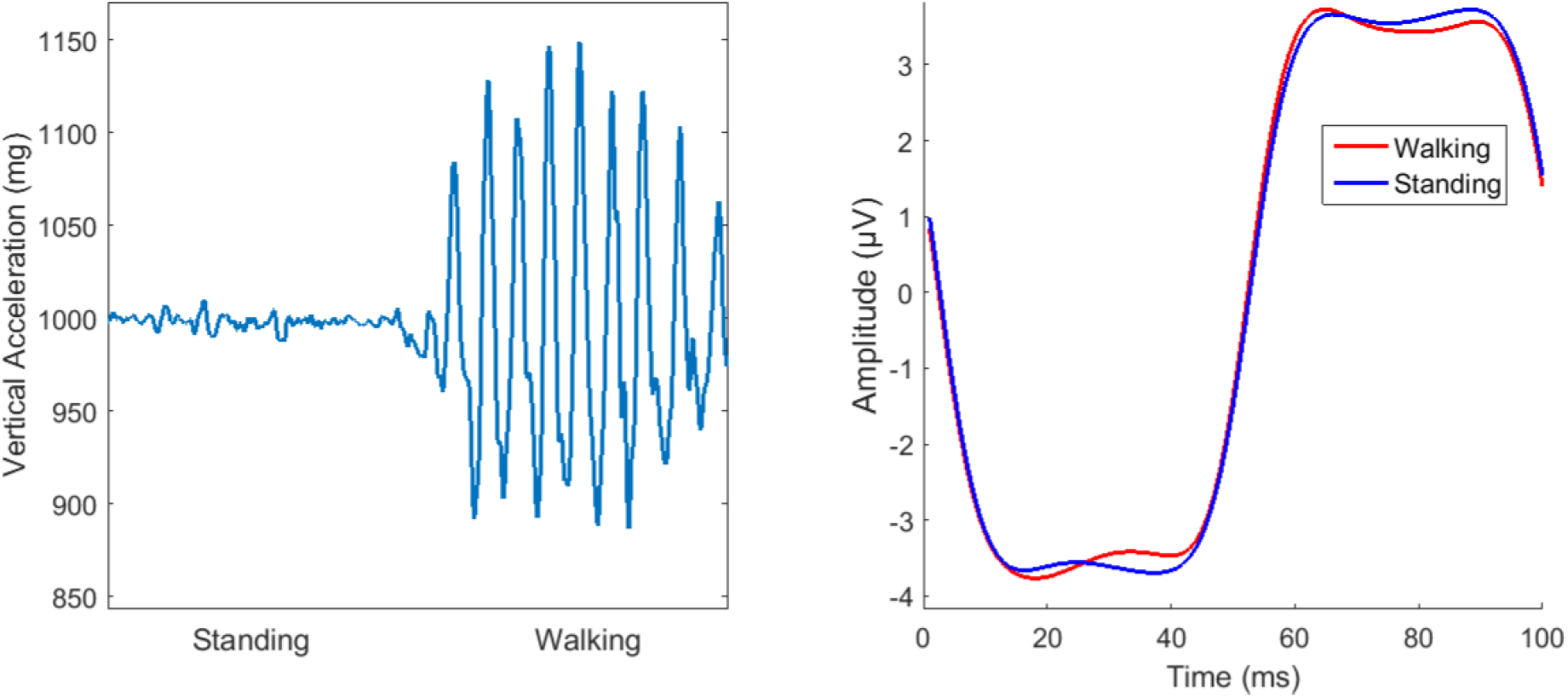
Left, example raw data trace from the vertical axis of the accelerometer attached to the head, showing motion during walking and standing. Right, example “simulated SSVEPs” from one participant (electrode P4) after averaging and filtering of the square wave signal. The high number of segments forming these averages results in these two traces being highly similar.

Before segmentation all data was high pass filtered to remove any slow drifts (1^st^ order Butterworth, 0.01 Hz high pass). SSVEPs were created by segmenting the data into 120 ms segments beginning 10 ms before each trigger, averaging all available segments, low pass filtering (2^nd^ order Butterworth, 40 Hz low pass) and then discarding the first and last 10 ms (20 ms in total) to remove any filter artefacts at the edges, leaving a 100 ms SSVEP beginning at the trigger, which corresponds to the darkening of the flickering glasses.

The “simulated SSVEPs” from the signal generator were created by segmenting 120 ms (one 100 ms cycle plus 10 ms either side) of data using the rising edge of the square-wave signal as a trigger, which was clearly visible in electrodes P7 and P8 where the signal was delivered. Trigger selection for walking and standing blocks, averaging and processing were done with exactly the same parameters as the real SSVEPs.

For each SSVEP the peak-to-peak amplitude was taken as the dependent variable. It is common in SSVEP experiments to perform a frequency transform (FFT) on a segment of the data to describe the amplitude of the evoked oscillation. However, here we chose not to do this for the main analysis: whilst the FFT is a highly convenient and useful method, recent research has brought into focus limitations which come with assuming neural oscillations are sinusoidal (Cole & Voytek, 2017). SSVEPs are often non-sinusoidal, complex waveforms, and comparing the waveform shape, and/or phase, can often provide information that might be missed with a simple analysis of power. Here the SSVEP was treated more like a traditional event related potential (ERP) and the peak to peak amplitude allowed the total size to be captured in a single number regardless of waveform shape. In addition, the correlation of phase/waveform shape between the SSVEPs in different conditions was analysed.

## Results

### Signal generator control condition

Visual inspection of the “simulated SSVEPs” from the signal generator control blocks showed very similar waveform and amplitude in the walking and standing conditions (Figure 2, right). This is a good qualitative indication that any additional noise added to the EEG signal during walking can be removed in the process of averaging with the number of segments used here (approximately 1800 - 2400). To quantify this, we took the average difference in peak-to-peak amplitude (standing minus walking) for the simulated SSVEPs: this was 0.07 μV, 0.27 μV, 0.22 μV and 0.15 μV for electrodes O1, O2, P3 and P4 respectively. This gives the average error when comparing an identical signal during walking and standing, introduced by the motion for a range of different walking speeds/gaits. This error is negligible compared to the sizes of the experimental effects below, demonstrating the efficacy of the method.

**Figure 3:**
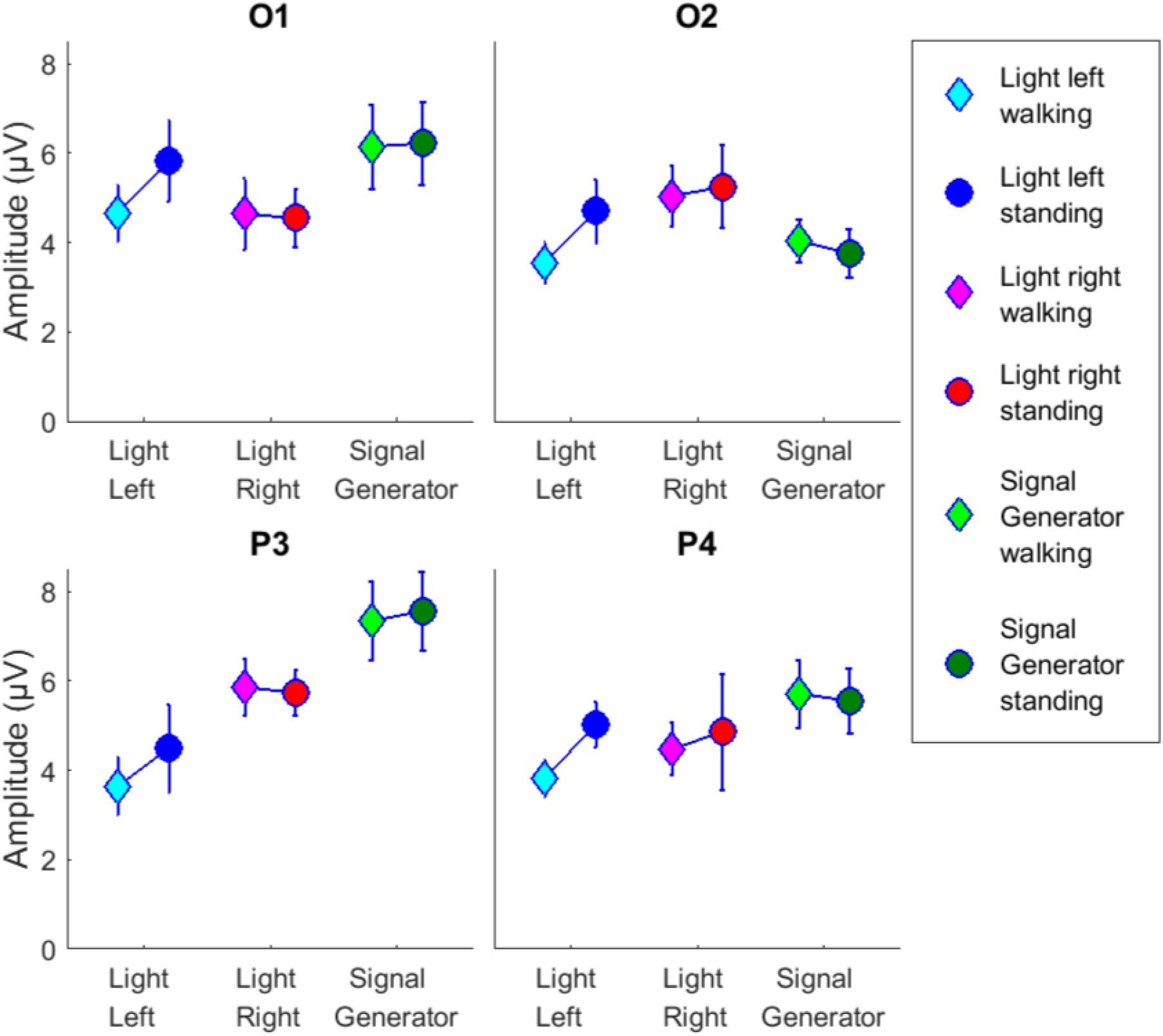
Mean SSVEP peak-to-peak amplitudes for each condition, error bars show the standard error of the mean. Data shown for left occipital (O1), right occipital (O2), left parietal (P3) and right parietal (P4) electrodes. Simulated SSVEPS from the signal generator (green) are within the same range as the real SSVEPs from walking (light colours) and standing (dark colours). Walking-Standing differences are shown in Figure 4.

### Walking vs. standing: SSVEP Amplitude

To test for differences during walking and standing in the real SSVEPs, the difference in peak-to-peak amplitudes was taken and compared to the difference between the simulated SSVEPs (figure 4). As data from 10 participants cannot be assumed to be normally distributed, non-parametric tests were applied; specifically, paired, two-sided, Wilcoxon signed rank tests between walking and standing conditions for each electrode. Bonferroni corrections were applied to control for multiple comparisons (4 electrodes x 2 conditions: adjusted alpha = 0.00625). Results show a significant difference between walking and standing only for left hemifield illumination and only for right hemisphere parietal electrode P4 (Z = 2.80, *p* = 0.005): SSVEPs during standing were significantly larger than walking. Inspection of the average SSVEP amplitudes for each condition from electrode P4 indicated that this effect is driven by a reduction in amplitude while walking only for the left hemifield illumination condition (Figure 3). Descriptively the SSVEPs were larger during standing with left hemifield illumination for all other electrodes but the variance was substantially greater and this did not reach significance after Bonferroni correction: O1 (Z = 1.5, *p* = 0.1), O2 (Z = −0.29, *p* = 0.016) and P3 (Z = 1.37, *p* = 0.17). For the right hemifield illumination condition there was no difference between standing and walking (all *p* values > 0.6). The mean difference between walking and standing for right hemifield illumination was near zero in all conditions: 0.08 μV, 0.18 μV, 0.13 μV and 0.39 μV for electrodes O1, O2, P3 and P4 respectively.

**Figure 4:**
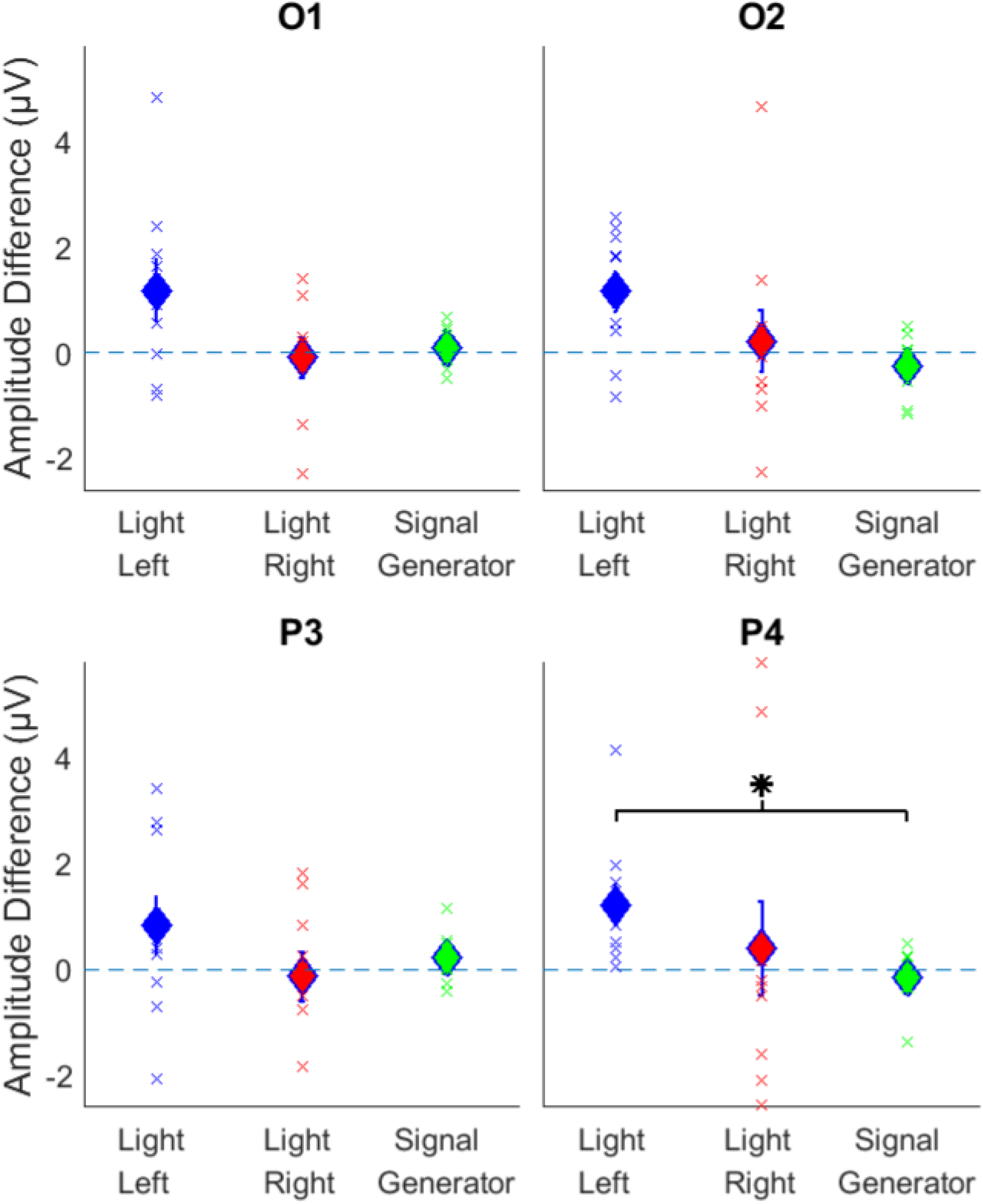
Difference in SSVEP peak-to-peak amplitude for each condition (standing minus walking); error bars show the standard error of the mean, dashed line indicates zero (i.e. no difference between conditions), crosses indicate individual participants’ mean differences. With left hemifield illumination the SSVEP amplitude difference at right parietal electrode P4 is significantly higher than with the signal generator, after Bonferroni correction.

### Walking vs. standing: Correlation

In addition to the amplitude of the SSVEPs the correlation between waveforms across conditions was also analysed. Visual inspection of the SSVEP waveforms indicated that the waveform shape and/or phase was highly different between left and right hemifield illumination conditions, but highly similar between walking and standing for any one hemifield being illuminated (Figure 5). To quantify this, the pairwise linear correlation coefficient was calculated as a measure of overall waveform similarity. Correlations were taken between the walking and standing SSVEPs and compared to the correlation between left and right hemifield illumination SSVEPs during standing for each participant (figure 6). Wilcoxon signed rank tests (Bonferroni corrected, 4 electrodes x 2 conditions, adjusted alpha = 0.00625) showed a significant difference between the correlation values of walking/standing and standing light-left/light-right for electrode P3 for both left and right hemifield illumination (Left, Z = −2.803 *p* = 0.005, Right Z = −2.803 p = 0.005). The same test for left hemisphere electrode O1 and standing/walking only reached marginal significance which did not survive Bonferroni correction (Z = −2.547 *p* = 0.010). Right Hemisphere electrodes (O2 and P4) did not show a significant difference (*p*‘s> 0.1), i.e. the walking/standing and left/right conditions both had highly similar waveform/phase.

**Figure 5:**
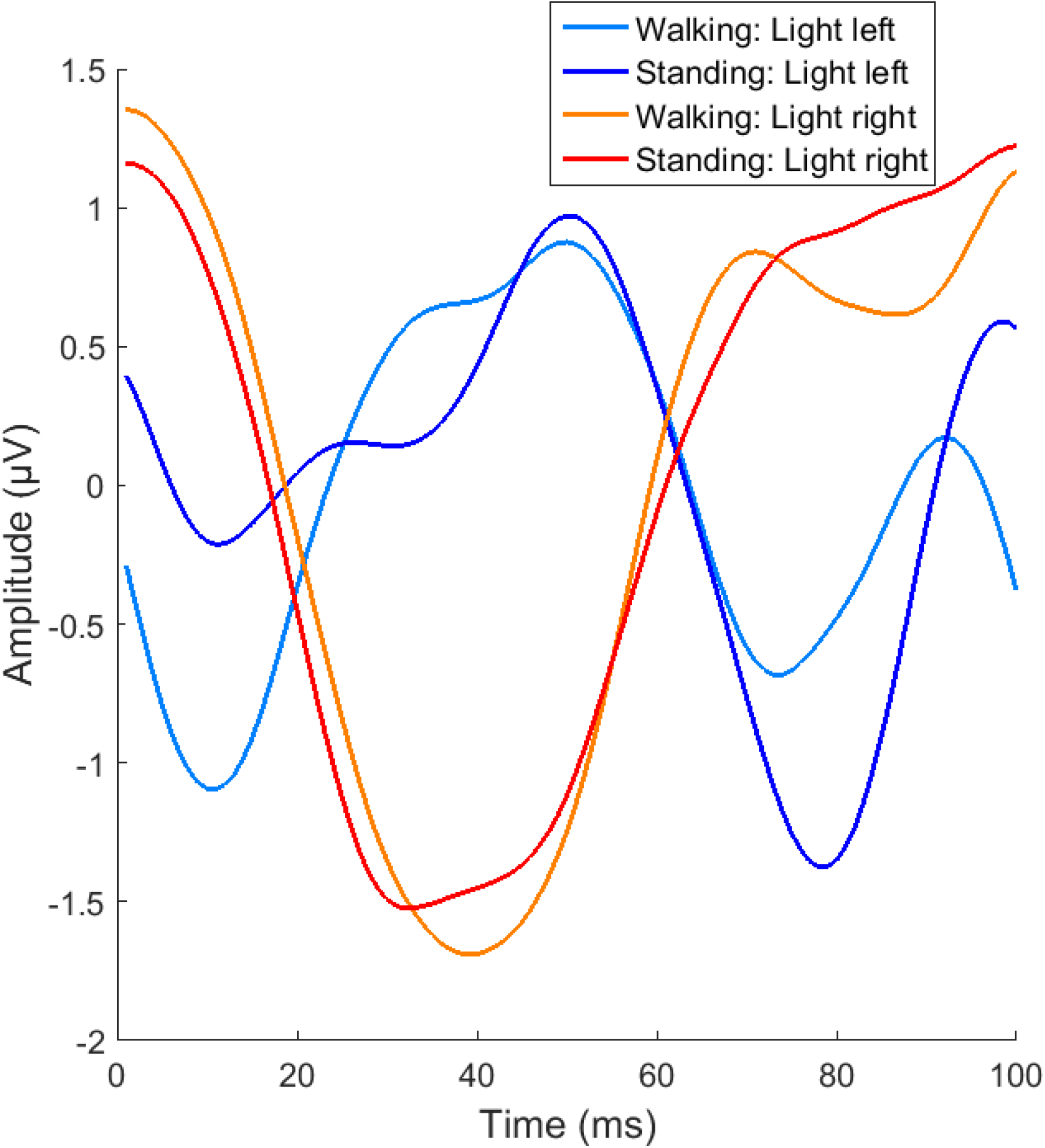
Example SSVEP waveforms from one participant (electrode P4). The phase and waveform shape is very different with left and right hemifield stimulation during standing, and is largely preserved when walking.

**Figure 6:**
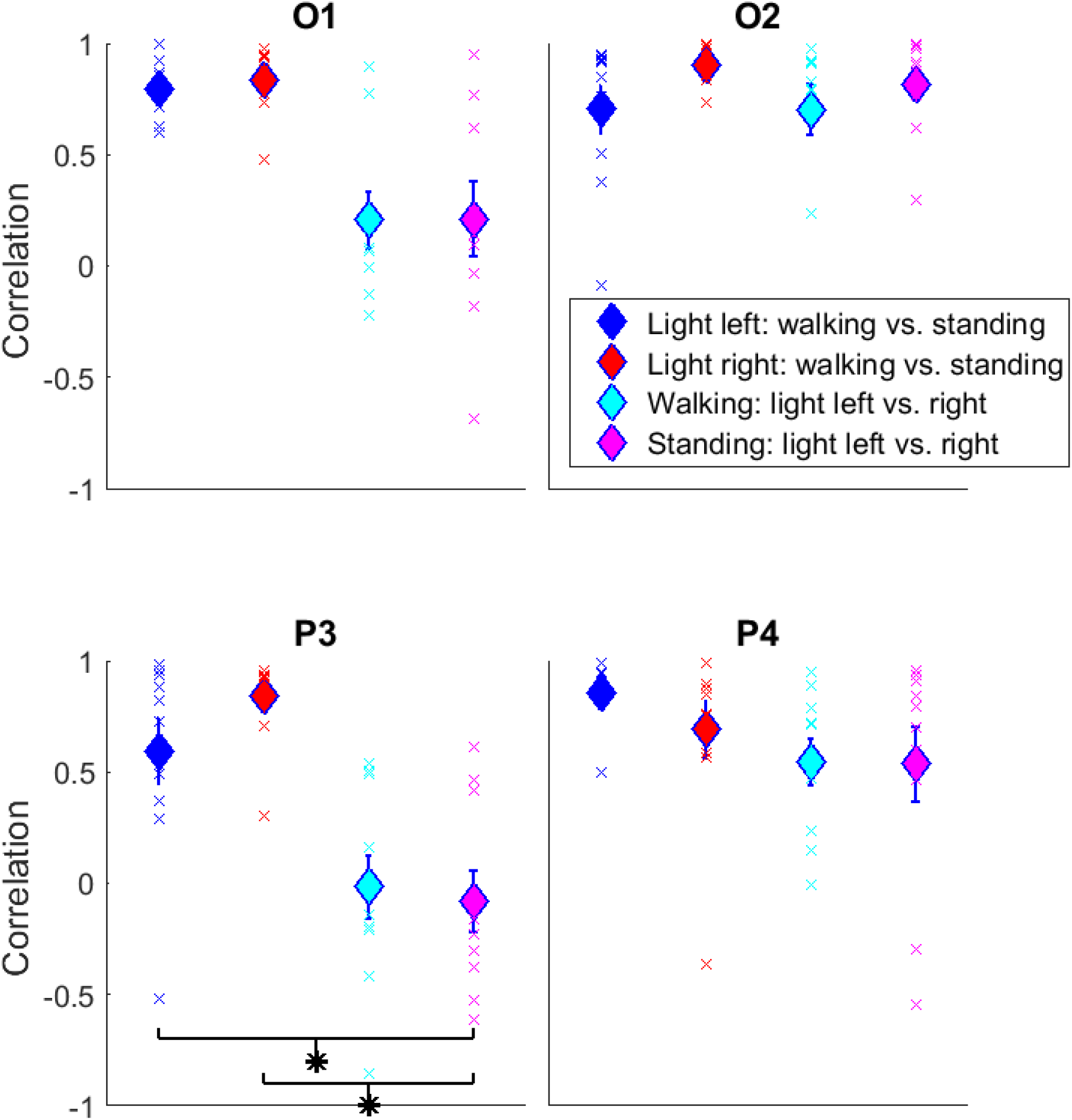
Mean correlation between the SSVEP waveforms across conditions. Error bars show the standard error of the mean, crosses show correlation values from individual participants. At left parietal electrode P3, the correlation between standing and walking is significantly stronger than the correlation between left and right hemifield illumination.

Correlation between walking and standing for the simulated SSVEPs was always above 0.95 indicating that phase/waveform shape is highly resilient to movement artefacts. For comparison, and to put this value in context, all available segments from each standing condition were randomly split into two groups to create two SSVEPs for P3 and P4 for each participant. We would expect these SSVEPs to be highly similar as they are taken from the same standing condition. The average correlations were 0.946 and 0.953 for light left, and 0.948 and 0.952 for light right; this is similar to the average correlation between walking and standing for the simulated SSVEPs, despite the substantial additional movement artefacts. This indicates that any very small residual differences in waveform between standing and walking, for the simulated SSVEPs, can be explained by the normal variance present within clean stable EEG, and is not appreciably affected by any additional movement artefacts produced during walking.

### Permutation tests on simulated SSVEPs

To be able to design future experiments with this method it is important to have an indication of how much additional noise is added to the SSVEPs during walking and how this varies with number of trials.

To quantify this factor, simulated SSVEPs were created from 1000 random subsets of the available segments during walking and each was compared to the simulated SSVEP in the standing condition with the full number of available segments, which is the closest available approximation of the “true” signal for each participant. The number of segments was initially set to 100, and increased in steps of 100 until 2000. For each step a random subset of the available segments of that number was used to create an SSVEP and this was compared to the “true” SSVEP in the standing condition. This was repeated 1000 times for each number of segments and the average difference was calculated, and then averaged across participants. The average difference between the walking SSVEPs and the “true” standing SSVEP, decreases exponentially with number of segments used (Figure 7, left) and converges towards zero, which would indicate no difference between standing and walking. This is quantified in how the standard deviation of simulated SSVEPs (across the 1000 repetitions) during walking converges on the value for standing after approximately 1500 segments are used (figure 7, right). Beyond this number of segments the standard deviation of the additional error during walking is essentially the same as for standing. 1500 segments from 10 Hz flicker would correspond to 150 seconds (2.5 minutes) of continuous walking, with the parameters used here.

**Figure 7:**
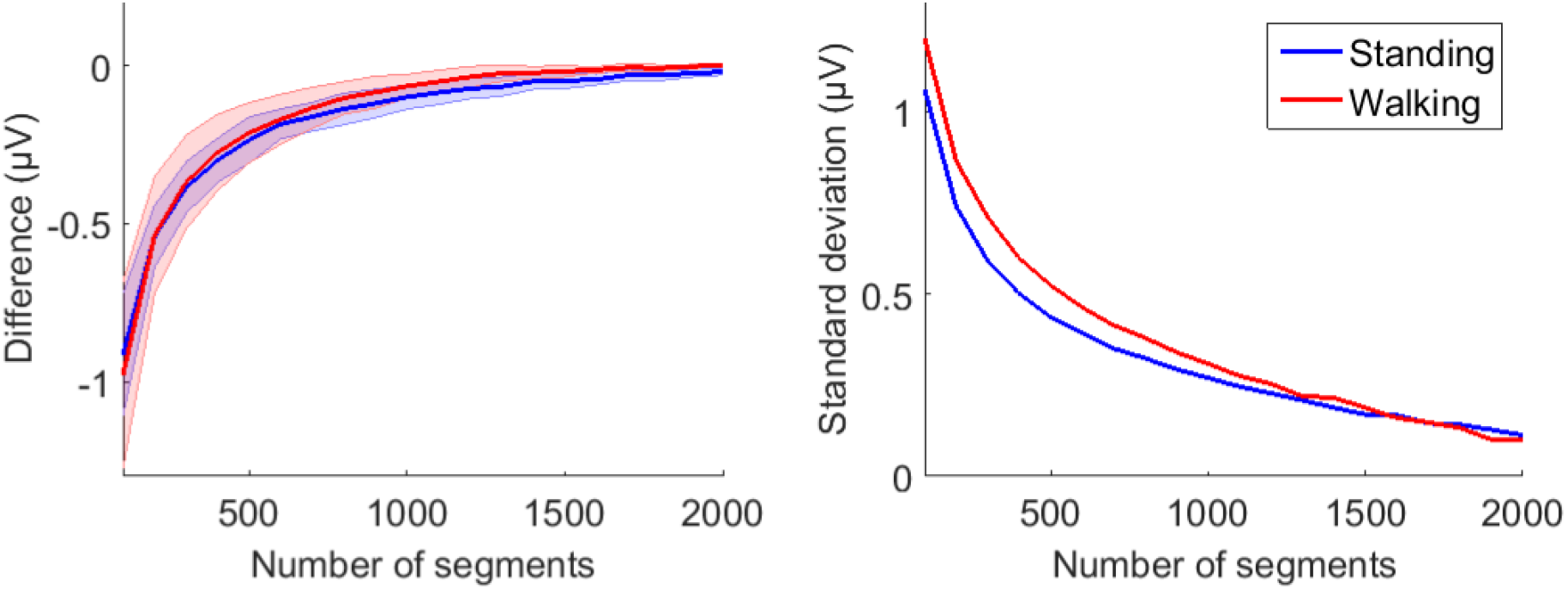
Left: Average difference between the “true” signal and 1000 simulated SSVEPs created from various numbers of segments, which decreases exponentially as the number of segments increases (shaded area shows standard error of the mean). Right: The mean standard deviation of the difference between the 1000 simulated SSVEPs when subtracted from the “true” signal. Data is from electrode P4 over right parietal cortex.

## Discussion

This study demonstrates that reliable Steady State Visual Evoked Potentials (SSVEPs) can be generated from viewing real world environments whilst walking. Thus, an effect of navigation on neural activity can be demonstrated. Additionally, key aspects of the visual scene can be differentiated within the mobile SSVEP; in this case the different illumination of the same visual scene, and with dissociable effects from locomotion.

The first main finding of this study is that SSVEPs can be recovered during walking with a meaningful signal to noise ratio, given enough trials, as demonstrated by a signal generator delivering an identical wave during walking and standing. The average difference between peak-to-peak amplitude of the simulated SSVEPs between walking and standing was less than 0.27 μV for all electrodes and the average correlation was above 0.95 for all electrodes and all individual participants. This correlation value is very similar to the average correlation between pairs of SSVEPs generated by random permutations from identical standing conditions (which would be expected to be highly similar as they are from the same condition without movement artefacts).

The second main finding is that SSVEP amplitude is reduced during walking, relative to standing, only for the left hemifield illumination condition, corresponding to the right hemisphere. This effect of navigation was significant for electrode P4, i.e. over right parietal cortex. Parietal cortex areas are known to be involved in encoding of space and time (Bremmer, Schlack, Duhamel, Graf, & Fink, 2001; Kaski et al., 2016). Although caution must be exercised in inferring anatomical localisation from EEG electrode position, a putative right lateralisation would be in agreement with the hypothesis that a unified perception of body position and self-motion requires the integration of information from the vestibular system with visual and somatosensory input and that this reflects the lateralization of the vestibular cortical system driven by the non-dominant hemisphere, i.e. the right hemisphere in right handers (Dieterich et al., 2003; Dieterich & Brandt, 2015, 2018). This vestibular dominance of the right hemisphere in right handers has been shown in several studies using both PET (Bense, S. Bartenstein, P. Lutz, S. Stephan, T. Schwaiger, M. Brandt, T. Dieterich, 2004; Dieterich et al., 2003) and MRI (Dieterich, Kirsch, & Brandt, 2017; Janzen et al., 2008; Schlindwein et al., 2008; Zu Eulenburg, Caspers, Roski, & Eickhoff, 2012). An interpretation of these results, which could be tested in future studies, is that the reduction in SSVEP amplitude seen here is due to the additional multisensory integration demands during walking engaging processing in right parietal cortex, for example to integrate visual and vestibular/somatosensory information, for perceptual stability and efficient navigation.

The third finding of this study is that it could be differentiated within the mobile SSVEPs whether visual stimuli had been presented to the left or right visual fields. This bolsters the potential applicability of this method in the future to tasks incorporating visual stimuli, and also serves as additional validation that a clean signal can be recovered despite movement. The pattern of this effect was very different to the locomotion effect, being present over the other hemisphere (left as opposed to right) and with a different measure (correlation as opposed to amplitude). Furthermore, the correlation between the waveform shape/phase of SSVEPs during walking and standing is most different, relative to a comparison of SSVEPs from visual stimuli presented to the left versus right visual hemifield, over left parietal cortex. One interpretation of this result would be that the waveforms from left and right hemifield illumination conditions are less correlated over left parietal cortex during standing, driving the effect seen here, indicating that the signal over left parietal cortex is more sensitive to the difference in left and right hemifield illumination of the room when standing. Further studies would be needed to test this hypothesis.

This method has wide-ranging potential applicability. Future research can test for differences in neural responses depending on a wide range of aspects of visual cognition, types of movement, and flicker frequencies. SSVEPs can be generated from flickering light up to 100 Hz (Herrmann, 2001) and a recent study (Herring, Herpers, Bergmann, & Jensen, 2019) has demonstrated SSVEPs of images flickering at gamma frequencies.

Direct comparison with the results presented here and other EEG studies should be approached with caution as there are certain key differences between viewing an actual scene and fixating on a computer monitor. For example the relative angle (vergence) of the eyes will be less when fixating on a wall 5 meters away than when viewing a screen close to the head. In the current study the vergence angle of the eyes will have increased as the participants walked towards the wall, but this factor was controlled for within the standing condition by the participants standing at increasing distances to the wall throughout the experiment.

The viewing of actual visual scenes, rather than precisely controlled visual stimuli on a computer monitor, may introduce additional inter-subject variability in the EEG responses, for example due to changes in the retinal image as a room is navigated (even during fixation of a cross, the retinal image of the cross would expand as it is approached, for example). Group averages of the SSVEP waveforms would result in very little signal, as the phase and waveform shapes of SSVEPs are very different across participants. The analysis strategy used in the current study was to compare SSVEPs of the same visual scene during walking and standing for each participant, which although different across participants, show a high degree of similarity within each participant (see Figure 6 for group correlations and Figure 5 for example data).

One potential source of variability in SSVEPs during walking, not present in the simulated SSVEPs, could be due to actual movement of the brain relative to the skull during walking, varying the distance between the cortex and the electrodes. However, any variability due to brain movement whilst walking would be the same in the left and right illumination conditions and could not account for the sensitivity to spatial location of visual stimulation.

## Conclusion

The highly controlled stimuli typically used in cognitive neuroscience experiments have many advantages, such as being reproducible, but at the expense of ecological validity. The assumption is that the neural correlates of pictures of visual scenes or virtual environments overlap meaningfully with the neural correlates of real environments into which we can physically move. The setup introduced here allows for probing the response of the visual system to virtually any visual scene, either real world or on a screen, at any frequency, i.e. not limited to multiples of the refresh rate of a computer monitor. This could provide important insights into the oscillatory dynamics of the visual cortex in response to various stimuli in different frequency bands during natural body movement.

